# Applying deconstructed sourdough communities and fermentation parameters to low-FODMAP wheat bread production

**DOI:** 10.1101/2024.12.16.628784

**Authors:** Anna Elizabeth Wittwer, Jiaying Amanda Li, Kate Howell

## Abstract

Despite being made with the same starting materials, standard and sourdough bread differ in terms of levels of fermentable oligosaccharides, disaccharides, monosaccharides, and polyols (FODMAPs). This study aims to assess the impact of a selected yeast-bacteria co-culture isolated from sourdough starters in conjunction with varying bread-making process parameters on bread FODMAP content and overall quality. A survey of the microbial composition of sourdough starters found that many contain *Maudiozyma humilis* and *Fructilactobacillus sanfranciscensis*. A selected pairing was used for small-scale bread-making, and dough fermentation and bread quality parameters were assessed. The sourdough microbes exerted distinctive effects on dough gassing capacity and FODMAP composition in comparison to baker’s yeast. To meet low-FODMAP cutoff values, the presence of a fermenting yeast and an extended dough fermentation time (> 6 h) were necessary. Sensory panellists preferred the sourdough co-culture bread fermented for 24 h over the 6 h treatment, and detected more complex aromas in the former. An extended, low-temperature dough fermentation regimen can be used to produce low-FODMAP bread with standard bread-making materials, and the use of sourdough fermentation microbes confers distinct textural and organoleptic properties.

## Introduction

Wheat bread is an important and widely consumed staple food of the Western diet. It is a good source of energy, dietary fibre, and micronutrients [1], while also being convenient and well-liked. However, certain wheat-based products are unsuitable for consumption by people with functional gastrointestinal disorders such as irritable bowel syndrome (IBS) because they are often rich in fermentable oligosaccharides, disaccharides, monosaccharides and polyols (FODMAPs).

Common examples of FODMAPs include fructans, α-galactooligosaccharides, lactose, fructose in excess of glucose, and polyols, and they are found in a broad range of food groups in addition to cereal-based products [2, 3]. These carbohydrates are not properly absorbed in the small intestine, leading to the production of gas and short-chain fatty acids in the large intestine by the gut microbiota. For this reason, many FODMAPs are also considered to be dietary fibres [4], but in IBS sufferers these properties lead to adverse gastrointestinal symptoms including abdominal pain, diarrhoea, flatulence, and bloating. Furthermore, certain FODMAPs attract water via osmotic activity, which can lead to luminal distention [5].

The major FODMAP present in wheat bread is fructans (**Figure 1**), which are β (2→1) and/or β (2→6) fructo-oligosaccharides (FOS) containing a single glucose moiety [6]. Fructan concentrations range from 0.6 to 2.9% in wholewheat flour [7], and 1.4 to 1.7% in white wheat flour [8]. The average degree of polymerisation (DP_av_) is 4, with a maximum of 14-19 [8]. Fructans derived from grain products are branched and categorised as graminan-type [2, 9]. Other FODMAPs present in wheat bread in smaller quantities can include raffinose, tri- and tetra-fructo-oligosaccharides (e.g., nystose, kestose), excess fructose, and mannitol [10].

**Figure 1.**
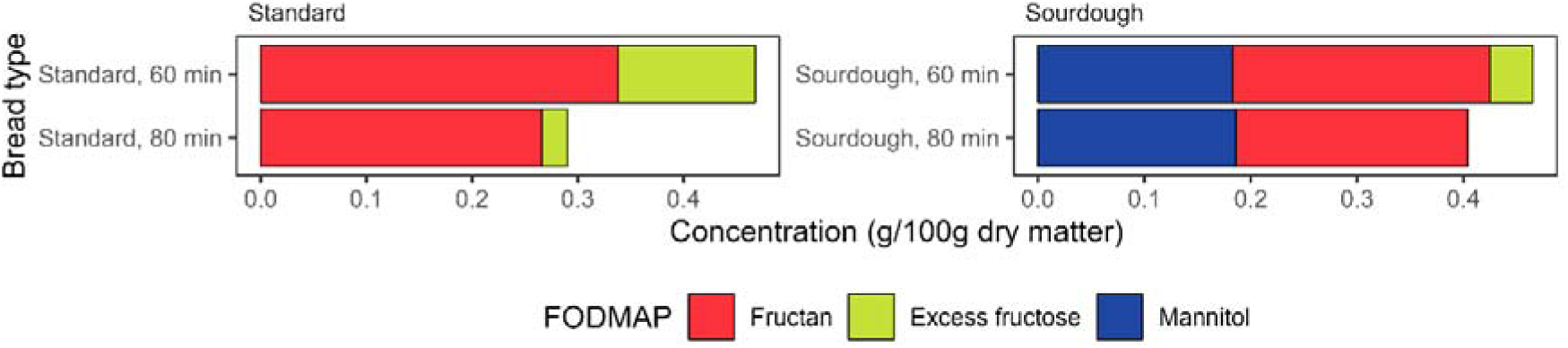
Key FODMAPs in standard and sourdough wheat bread fermented for 60 or 80 minutes. Bars represent mean values. Data from Schmidt and Sciurba [6].

A diet low in FODMAPs is a proven therapeutic measure to reduce or eliminate IBS symptoms in those with functional gastro-intestinal disorders [11], but the exclusion of wheat bread from the diet is not ideal due to its staple food status and the non-FODMAP fibres and micronutrients it provides. Bread can be considered low-FODMAP, however, if in a serving of bread (50 g) the following cutoff values are not exceeded: 0.3 g of oligosaccharides, 0.15 g of excess fructose, 0.4 g of polyols, and a total of 0.5 g of FODMAPs excluding lactose [3]. Current low-FODMAP breadmaking focuses on either enzymatic pre-treatment of flour to remove FODMAPs, or the use of grains that are naturally low in FODMAPs, such as spelt [10]. While both options are viable, they are relatively expensive and are not as broadly accessible as conventional white wheat bread.

Conventional white wheat bread produced at commercial scales is typically leavened with baker’s yeast (industrial strains of *Saccharomyces cerevisiae*) for a short period of time at approx. 30°C [12]. Despite the fact that baker’s yeast produces invertase, an enzyme that can effect fructan hydrolysis [13], FODMAP levels in conventional, rapidly-fermented wheat bread often exceed low-FODMAP thresholds, mainly because of high fructan content (**Figure 1**).

Sourdough bread, on the other hand, while made with the same ingredients as standard bread, has a distinct and often improved FODMAP composition. Namely, its fructan content is lower, whereas levels of excess fructose and mannitol may be enriched [6]. Sourdough bread is traditionally leavened with a type I sourdough starter. This is a spontaneously-fermented mixture of flour and water that via a process of regular re-feeding with fresh ingredients (termed ‘backslopping’) develops a community of often non-conventional yeasts and bacteria [14, 15]. A mature sourdough typically contains one or two yeast species and one-several bacterial species [16], which may interact with one another by various means. One of the most common sourdough yeasts is *Maudiozyma humilis* (formerly *Kazachstania* or *Candida humilis*), an ascomycetous yeast that occupies a phylogenetically close position relative to the *Saccharomyces* genus [17–19]. This yeast is commonly isolated from sourdough starters that also contain *Fructilactobacillus sanfranciscensis*, an obligately heterofermentative lactic acid bacterium (LAB) whose primary ecological niche is sourdough starters [20, 21]. In addition to its varied microbial profile compared to conventional bread, sourdough bread is often made according to different process parameters. Depending on the style of sourdough production, dough proofing or fermentation can be carried out over approx. 24 h and can include several propagation steps [22].

It is already well-understood that extending dough proofing times leads to increased degradation of fructans in bread dough [6, 23]. However, the effects of proofing temperature on this process and its interaction with proofing time are less well-characterised. Furthermore, while some yeasts, such as *Saccharomyces cerevisiae*, *Kluyveromyces marxianus*, and *Lachancea fermentati* have been investigated for their applicability to low-FODMAP breadmaking [1, 24], there remains a lack of information about the potential of common sourdough microbes in this area, given the low-fructan status of sourdough bread.

Therefore, the aim of this study was to investigate how the functional metabolic properties of a common sourdough yeast-bacteria pairing and the dough proofing time-temperature regime typical of sourdough breadmaking affect bread quality and FODMAP content. An accompanying aim was to draw comparisons between conventional and sourdough bread conditions.

## Materials and Methods

### Growth media preparation

For yeast strain isolation, two media were prepared. Wallerstein Nutrient (WLN) agar was prepared according to the manufacturer’s instructions (Oxoid). Yeast-peptone-dextrose (YPD) agar was composed of 10 g/L yeast extract (Oxoid), 20 g/L bacteriological peptone (Oxoid), 20 g/L anhydrous D-glucose (Merck), and 20 g/L w/v bacteriological grade agar (Ajax). When used for strain isolation and viable colony enumeration, yeast growth media were supplemented with 100 μg/mL ampicillin and 30 μg/mL chloramphenicol to prevent bacterial growth.

For bacterial strain isolation, de Man, Rogosa & Sharpe (MRS) agar was prepared according to the manufacturer’s specifications (Oxoid). Isolation and enumeration media were supplemented with 10 μg/mL cycloheximide to prevent fungal growth.

Antibiotic solutions were added aseptically to just-autoclaved media at approx. 40-50 °C.

### Strain isolation and identification

Nine sourdough starters were donated to this project. Codes, suburbs and isolates from each bakery are given in **Table 1**.

**Table 1.** *Maudiozyma humilis* and *Fructilactobacillus sanfranciscensis* isolated from sourdough starters. Cells shaded grey indicate starters that contain both species.

| Leaven | Origin | Code | Species | Top BLAST identity (%) |
| --- | --- | --- | --- | --- |
| Os | Ovens St Bakery, Brunswick VIC | OsY1 | *Maudiozyma humilis* | 98.9 |
| Osr | Ovens St Bakery, Brunswick VIC | OsrY6 | *Maudiozyma humilis* | 99.3 |
| Os | Ovens St Bakery, Brunswick VIC | OsY5 | *Maudiozyma humilis* | 99.4 |
| Osr | Ovens St Bakery, Brunswick VIC | OsrY1 | *Maudiozyma humilis* | 99.5 |
| Nrk | Natimuk, VIC | NrkY1 | *Maudiozyma humilis* | 99.7 |
| Nrk | Natimuk, VIC | NrkB1 | *Fructilactobacillus sanfranciscensis* | 100.0 |
| Os | Ovens St Bakery, Brunswick VIC | OsB1 | *Fructilactobacillus sanfranciscensis* | 100.0 |
| Os | Ovens St Bakery, Brunswick VIC | OsB2 | *Fructilactobacillus sanfranciscensis* | 100.0 |
| Osr | Ovens St Bakery, Brunswick VIC | OsrB1 | *Fructilactobacillus sanfranciscensis* | 100.0 |
| Lbr | Loafer Bread Bakery, Fitzroy North VIC | LbrY1 | *Maudiozyma humilis* | 98.0 |
| Fb | Falco Bakery, Collingwood VIC | FbY1 | *Maudiozyma humilis* | 99.7 |
| Lb | Loafer Bread Bakery, Fitzroy North VIC | LbY2 | *Maudiozyma humilis* | 99.8 |
| Gw | Great Western Granary, Great Western VIC | GwB1 | *Fructilactobacillus sanfranciscensis* | 100.0 |
| Kb | Ket Baker, Wallington VIC | KbB1 | *Fructilactobacillus sanfranciscensis* | 100.0 |
| Zb | Zelda Bakery, Ripponlea VIC | ZbY4 | *Maudiozyma humilis* | 100.0 |

During sampling, care was taken to avoid contamination such that only microbes present in the sourdough starter itself or the ‘house microbiota’ (organisms associated with each isolate’s respective bakers and bakery environment) of the bakery would be collected [25]. Bakers were asked to collect a portion of their sourdough starter(s) in sterile tubes, or to provide a sample in their own container. Samples were stored at room temperature prior to analysis and cultured 24-72 h after the last starter refreshment.

Sourdough starters were first serially diluted by adding approx. 1 g starter to 9 mL 0.1% w/v peptone water. Suspensions were vortexed thoroughly between each dilution step. Serial dilutions were carried out until a factor of 10^-6^ was reached. Aliquots of 100 μL of each 10^-3^-10^-6^ suspension were spread-plated on antibiotic-supplemented WLN and YPD agar to isolate yeasts, and antibiotic-supplemented MRS agar to isolate LAB.

Plates were incubated in aerobic conditions at 28 °C for 2-3 days. Plates with 30-300 countable colonies present were selected for sub-culturing. Well-isolated colonies were selected on the basis of distinct morphological characteristics. Three iterations of streak plating were performed to isolate pure colonies. After the third iteration, each isolate was given an identification code to reflect its origin, and glycerol stocks were stored at -80 °C.

#### Yeast isolate identification

Genomic DNA was extracted from pure yeast colonies with the MasterPure™ Yeast DNA Purification Kit (Epicentre®, Madison, WI) according to the manufacturer’s instructions. PCR amplifications of the extracted DNA were carried out using the KAPA HiFi HotStart PCR Kit (Kapa Biosystems, Wilmington, MA). The reaction used the primers NL-1 and NL-4 to target the D1/D2 region of the large subunit ribosomal DNA as described in Kurtzman & Robnett [26]. Amplicons were submitted to the Australian Genome Research Facility (AGRF) for sequencing. Raw sequences were trimmed, forward and reverse sequences aligned, and the alignments searched against the NCBI nucleotide database in Geneious Prime 2024.0. In the event of an ambiguous or improbable top result, YeastIP [27], a curated sequence database, was used to identify isolates.

#### Bacterial isolate identification

Pure bacterial colonies were prepared for identification in Applied Biosystems™ PrepMan® Ultra Sample Preparation Reagent (Thermo Fisher Scientific). Attenuated samples were submitted to the AGRF for taxonomic assignations using the 16S rRNA gene.

### Strain selection for dough and bread analyses

One sourdough-derived *Maudiozyma humilis* strain was selected for small-scale breadmaking trials based on the following criteria: a) originated from a sourdough starter in which a *Fructilactobacillus sanfranciscensis* isolate was also present and b) demonstrated acceptable levels of growth on standard growth carbohydrates and fructan-like carbohydrates.

A defined medium (Yeast Nitrogen Base with amino acids (YNB) Sigma) supplemented individually with 0.5% w/v glucose, fructose, small FOS (DP 2-8), or large FOS (DP > 60) was prepared to test yeast isolate growth on different carbon sources. Small FOS and large FOS (inulin) were purchased from Megazyme (Bray, Ireland)

Yeasts were precultured twice in aliquots of 5 mL of a starvation medium based on the composition of YPD that contained 5% of the standard concentration of glucose (1% w/v yeast extract, 2% w/v peptone, 0.1% w/v glucose). Cell suspension turbidity was standardised to an optical density measurement at 600nm (OD_600_) of 0.1 in 0.1% peptone. To each well of a U-bottomed 96-well microtitre plate, 198 μL supplemented YNB broth was added. Aliquots of 2 μL of the yeast cell suspensions were added, and 2 μL 0.1% peptone was used for the blanks. Plates were incubated for 24 h at 28 °C, and cell density was measured with a plate reader (Multiskan Go®, ThermoFisher) every hour at 600 nm. Kinetic parameters used to assess yeast growth performance were obtained with the *growthcurver* package [28].

### Dough and bread analyses

#### Inoculum preparation

Yeasts and bacteria were maintained on YPD and MRS agar, respectively. A fresh plate was prepared 2-5 days prior to breadmaking to ensure maximum live cells. *M. humilis* OsrY1 and the baker’s yeast were precultured in 30mL YPD broth at 28 °C at 220 rpm for 18 h. *F. sanfranciscensis* OsrB1 was precultured in 30mL MRS broth at 29 °C at 18% CO_2_ for 72 h. Prior to baking, precultures were centrifuged at 4000 rpm/1789 rcf for 5 min and the growth medium was removed.

To inoculate dough with baker’s yeast in the same manner as the sourdough-derived microbes, a commercially available baker’s yeast was purchased and cultured in YPD broth. The baker’s yeast was maintained on YPD and stored in 80% v/v glycerol.

#### Dough preparation

Bread dough was prepared in small domestic breadmaking machines (BM2500 Compact Bakehouse, Sunbeam, NSW, Australia). Each batch of dough was composed of 500 g flour, 7.5 g table salt, 5 g canola oil, and 315 g water. Pre-cultured cells were added as shown in **Table 2** with a sterile transfer pipette. White wheat all-purpose baker’s flour containing 11.5% protein and 3.0% dietary fibre was used (Wallaby T55 variety, Laucke, South Australia). Doughs were mixed for 5 min on setting 9, then the machines were switched off.

**Table 2.** Dough fermentation microbe treatments applied to samples.

| Treatment, code | Microbe(s) | Inoculum mass (g) | CFU/g dough |
| --- | --- | --- | --- |
| Unstarted dough, None | None | 0 | 0 |
| Baker's yeast, BY | *Saccharomyces cerevisiae* BY | $0.6 \pm 0.05$ | $2.4 \times 10^5$ |
| Sourdough yeast, OsrY1 | *Maudiozyma humilis* OsrY1 | $0.6 \pm 0.05$ | $2.4 \times 10^5$ |
| Sourdough bacterium, OsrB1 | *Fructilactobacillus sanfranciscensis* OsrB1 | $0.6 \pm 0.05$ | $8.6 \times 10^6$ |
| Sourdough co-culture, OsrY1 + OsrB1 | *M. humilis* OsrY1 + *F. sanfranciscensis* OsrB1 | $1.2 \pm 0.1$ | $2.4 \times 10^5$ yeast + $8.6 \times 10^6$ bacteria |

#### Dough characterisation

##### Enumeration of yeast and LAB

Quantities of 0.5 ± 0.05 g bread dough were added incubated at 25 or 32°C. At 0, 2, 6, and 24 h proofing, aliquots of 4.5 mL 0.1% w/v peptone water were added. Tubes were vigorously shaken and vortexed to create a 10^-1^ dilution, then the dough slurry was serially diluted to 10^-5^. Aliquots of 75 μL of the 10^-3^-10^-5^ dilutions were spread plated on antibiotic-supplemented MRS and YPD agar. The MRS plates were incubated at 27°C under 18% CO_2_ and the YPD plates were incubated under aerobic conditions at 28°C for 4 days. Plates containing colonies within the countable range (30-300) were selected for enumeration.

##### Dough pH and temperature

A solid-state food and dairy pH meter HI199161 (Hanna Instruments) was used to measure dough pH and temperature at each proofing time point. Three separate measures were obtained at each time point by placing the pH probe in three different locations within the dough sample.

##### Dough gassing capacity

Immediately after mixing, portions of 100 g dough were added to 500 mL Schott bottles and sealed with gas pressure modules (ANKOM Technology, Macedon, NY). Bottles were kept in incubators set to 25 or 32°C, and pressure measurements were taken every 15 s over 24 h.

##### Dough FODMAP quantification

Quantities of approx. 5 g dough were sampled at 0, 6, and 24 h of fermentation. The 2 h timepoint was omitted from analysis due to unacceptable bread technical qualities (i.e. insufficient gas production for leavening), but the 0 h was analysed as a baseline. Dough samples were freeze-dried for 72 h (Biobase freeze dryer BK-FD10P). Dough was then crushed to a powder with a metal spatula, and FODMAPs were extracted using a hot water method. For fructan analyses, 0.1 g freeze-dried, powdered dough was added to 5 mL deionised water and incubated in boiling water for 15 min. For the other FODMAP and dough enzyme analyses, 0.1 g lyophilised dough was added to 1 mL deionised water and incubated for 10 minutes at 80°C in a heating block. Fructan, mannitol, and excess fructose were quantified with UV-based methods using kits from K-FRUC, K-MANOL, and K-FRUGL kits respectively (Megazyme).

#### Bread preparation and characterisation

Quantities of 325 g dough were portioned into lightly oiled containers, covered with cling film and left to prove for 2, 6, or 24 hours in incubators set to 25 or 32°C. One hour before the end of the set proofing time, doughs were knocked back and shaped into small loaf tins (14.6 x 7.62 cm, Wilton). For all doughs, the final hour of proving was carried out at 25°C to ensure consistent dough temperature prior to baking. Loaves were baked in a Convotherm 4 easyDial 6.10 ES oven (Elfging, Germany) for 30 min at 180°C (fan 4, Crisp & tasty 3, Bake pro 3). After baking, they were cooled, and a serrated knife was used to make slices of approx. 2 cm thick. The maximum height of the middle three slices from each loaf was measured.

Two slices from each loaf were photographed with an iPhone 13 using default camera settings in a light box (Photosimile 50; Ortery, Irvine, CA) for digital image analysis. Photographs were cropped to show only the largest area of regular crumb patterns. They were then analysed with MATLAB® ver. R2023b (Mathworks Inc., Matick, MA) to output various parameters, including void fraction, mean cell area (mm^2^), small cell density (per cm^2^), and large cell density (per cm^2^).

#### Bread sensory analysis

An untrained sensory consumer panel was used to test palatability and acceptability of low-FODMAP wheat breads. Four samples were selected based on a) low-FODMAP status and b) acceptable bread technical quality (Table 3).

**Table 3.**
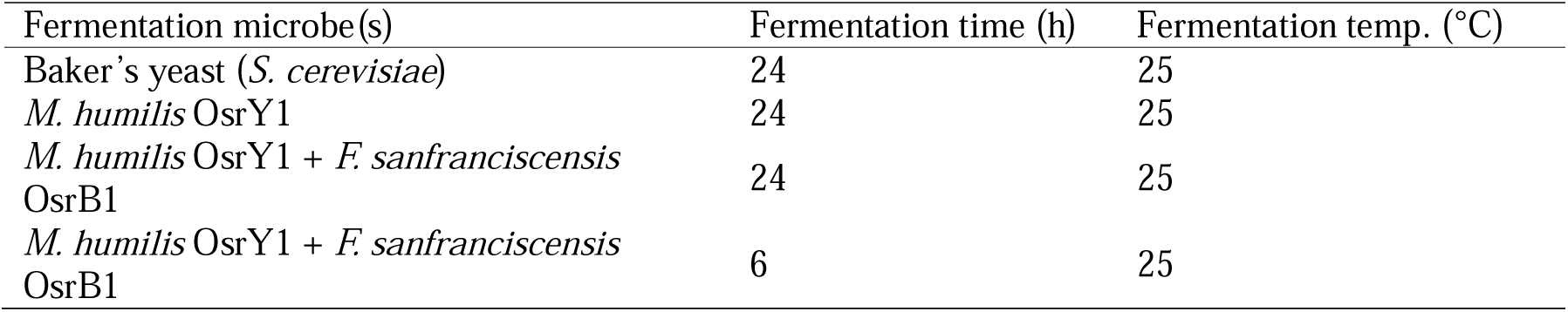
Low-FODMAP bread samples chosen for sensory analysis.

Participants (*n* = 56) aged 18-50 attended four sensory panel sessions held over a 5-day period. One week prior to testing, bread loaves were baked, cut into 1-cm-thick slices, and frozen. Samples were defrosted the night before each session. Participants were asked to examine and taste each bread sample one at a time, labelled with a random 3-digit code. The order in which the four samples were given was randomised for each participant. They were asked to score each sample based on how it looked, smelled, and tasted using a 7-point semi-structured scale with described extremes. Approval for this experiment was given by the University of Melbourne Human Research Ethics Committee (HREC 2024-30068-56367-3).

### Statistical analyses

All experiments were carried out in triplicate, and statistical analyses and data visualisations were completed in R [29]. Group means for all data except the bread sensory data were compared with the Kruskal-Wallis nonparametric test, and *post-hoc* pairwise comparisons were made with the Dunn’s test with Bonferroni’s correction applied. Sensory data scores were compared with one-way ANOVA, and pairwise comparisons conducted with Tukey’s HSD test. Statistical significance was considered at *p* < 0.05.

## Results

### Yeasts and bacteria from sourdough starters

Based on LSU rDNA gene sequencing results, nine *Maudiozyma humilis* strains were isolated from the sourdough starters studied (**Table 1**). Pairwise identity for yeasts ranged from 98 to 99.8% and for bacteria was 100%. Bacteria were identified using the 16S rRNA gene, and the main species identified was *Fructilactobacillus sanfranciscensis*.

Since *M. humilis* and *F. sanfranciscensis* are a frequently co-occurring pair in sourdough starters [15], these species were selected for low-FODMAP bread-making trials. In order to maximise potential FODMAP degradation, we focused on assessing the functional properties of the *M. humilis* strains, as *F. sanfranciscensis* is not among the group of LAB that produce FODMAP-degrading enzymes such as fructanase [30]. To select the *M. humilis* strain most capable of bread FODMAP reduction and best suited to bread-making, growth was measured in defined media individually supplemented with carbohydrates. Strains with growth kinetic parameters are listed in **Table 4**.

**Table 4.**
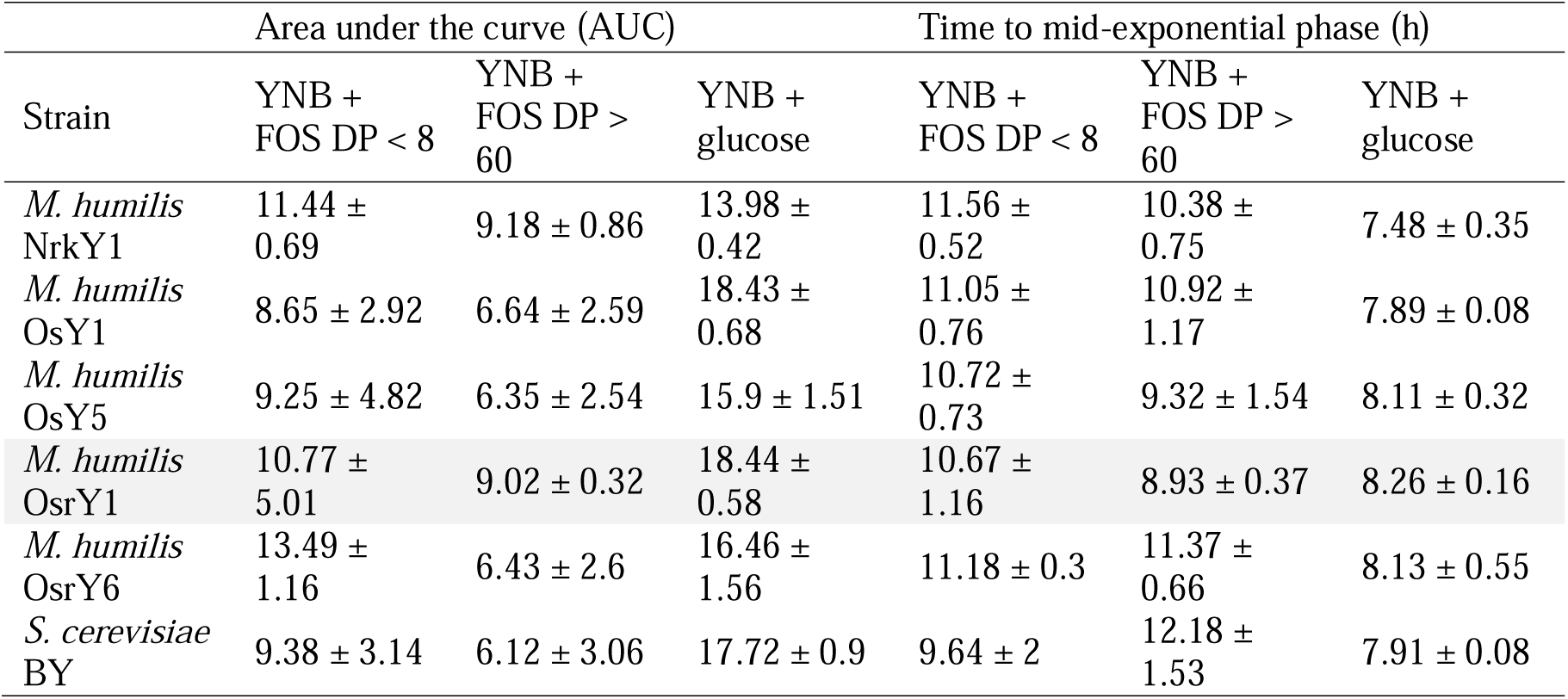
Growth kinetic parameters of *Maudiozyma humilis* strains. Growth performance of strains was assessed in yeast nitrogen base (YNB), a defined medium individually supplemented with fructo-oligosaccharides (FOS) with a degree of polymerisation (DP) of 2-8, FOS DP > 60, or glucose. Values shown are mean ± SD.

*M. humilis* OsrY1 was selected for small-scale breadmaking trials because its growth intensity and speed in defined media were either equivalent or superior to those of the other strains, particularly on media supplemented with large FOS (DP > 60, **Table 4**). *F. sanfranciscensis* OsrB1 isolated from the same starter (**Table 1**) was selected for co-culture and bacterial mono-culture tests.

### Microbial growth kinetics

Bread dough was prepared with the following five microbe conditions: baker’s yeast, *M. humilis* OsrY1, *F. sanfranciscensis* OsrB1, *M. humilis* OsrY1 + *F. sanfranciscensis* OsrB1, and no inoculum (unstarted dough). Doughs were fermented in proofing ovens set to either 25 or 32°C.

Internal dough temperature stabilised by 2 h fermentation and did not change thereafter. The mean internal temperature of doughs fermented at 32°C and 25°C, regardless of microbial treatment, was 29°C and 24°C, respectively (**Table S2**).

Growth and persistence of yeasts and LAB in the dough substrate was investigated through plate count-based enumeration (**Figure 2**). For all four inoculated treatments, colony-forming unit (CFU) counts increased significantly over the course of the total 24 h fermentation time. This increase was only statistically significant for the co-culture dough when fermented at 32°C, however. In the three mono-culture doughs inoculated with only one strain, no contaminating yeast or bacteria were detected. By 24 h, the baker’s yeast reference dough did not have significantly higher yeast counts than the *M. humilis* OsrY1 mono- and co-culture doughs.

**Figure 2.**
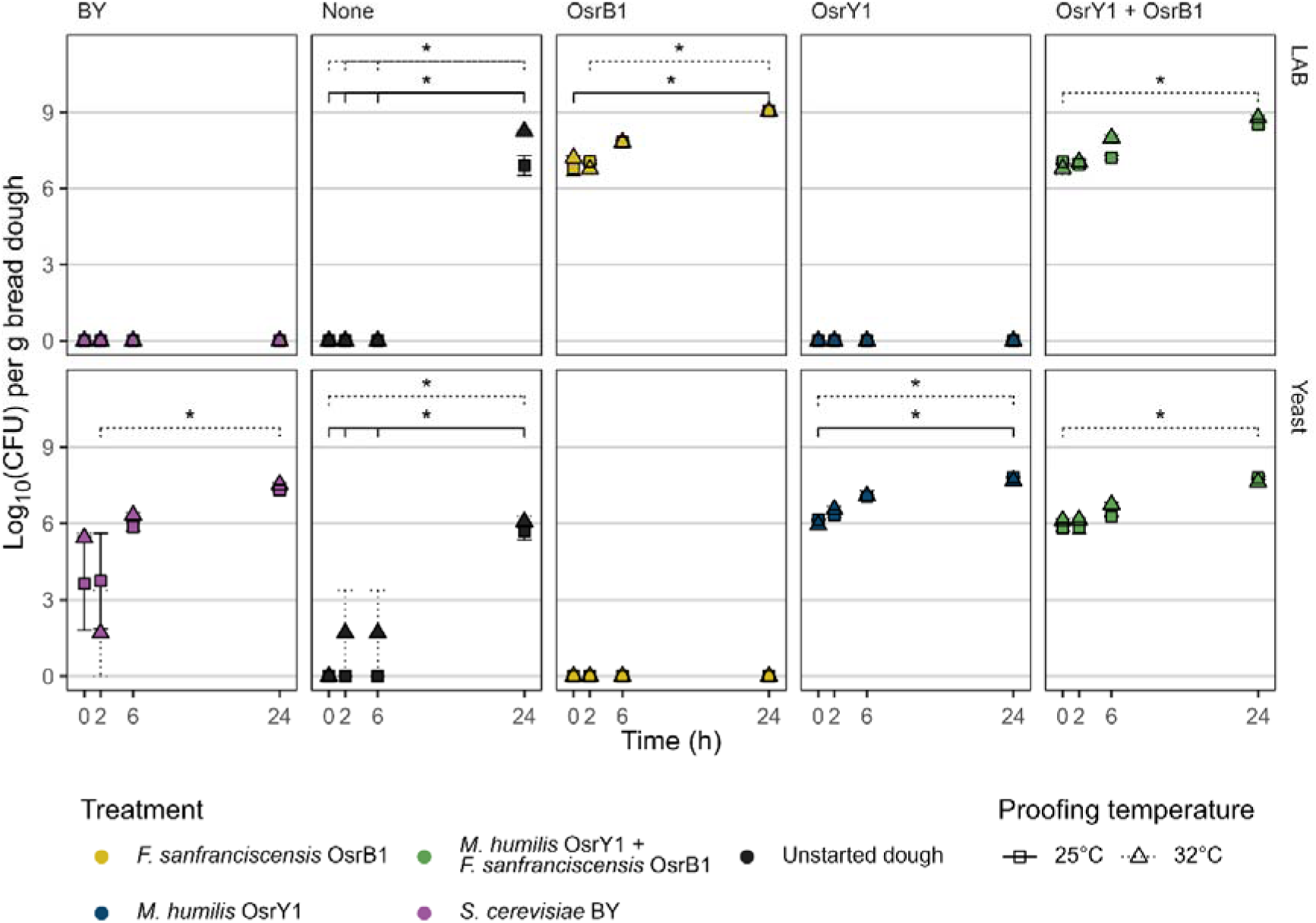
Changes in viable yeast and LAB counts in dough fermented at 25 and 32°C over time. Dough samples were serially diluted in 0.1% w/v peptone, and yeast and LAB were enumerated on antibiotic-supplemented YPD and MRS agar plates respectively. Log_10_-transformed data presented as mean (*n* = 3) colony-forming units (CFU) ± SE, * represents *p* < 0.05.

The highest CFU/g dough values were found in the dough inoculated solely with *F. sanfranciscensis* OsrB1. Proofing temperature was found to have a significant effect on growth. At 6 h, LAB counts were significantly higher in the co-culture dough fermented at 32°C than those in the 25°C co-culture dough. No significant differences between LAB counts in the *F. sanfranciscensis* mono-culture dough proofed at 25 and 32°C were detected. Conversely, for both treatments containing *M. humilis* OsrY1, yeast counts at 24 h of fermentation were higher at 25°C than at 32°C. In the unstarted dough treatment, to which no microbes were added during the dough mixing stage, counts of LAB and yeast increased markedly by 24 h, and this increase was more pronounced at 32°C than at 25°C.

Dough pH was measured with a solid-state meter throughout the fermentation process. Over the 24-hour measurement period, the pH of all doughs, regardless of inoculum or proofing temperature, decreased (**Table S1**). pH of the *M. humilis* mono- and co-culture doughs decreased significantly from 2 to 24 h at both temperatures. In all dough fermentation treatments, the 24-hour samples had lower pH values when fermented at 32°C than when fermented at 25°C.

### Fermentation dynamics

An ANKOM gas production system was used to measure each treatment’s dough leavening capability in small quantities over 24 h. Area under the curve (AUC) of each treatment-temperature combination was calculated to quantify overall gas production.

Dough cumulative pressure output was affected by microbial treatment, proofing temperature, and individual treatment-temperature combinations. The treatments can be grouped into high- and low-gassing capacity status according to whether they contain a yeast or not; the latter group’s mean AUC is < 50 (**Figure 3B**). Among these groups, the treatments that demonstrated the highest and lowest gassing capacity were baker’s yeast fermented at 32°C and unstarted dough at 25°C. Overall gas production at either fermentation temperature did not significantly differ between any of the yeasted treatments. At 2, 6, and 24 h, dough leavening ability was significantly improved by increased fermentation temperature (**Figure 3C**).

**Figure 3.**
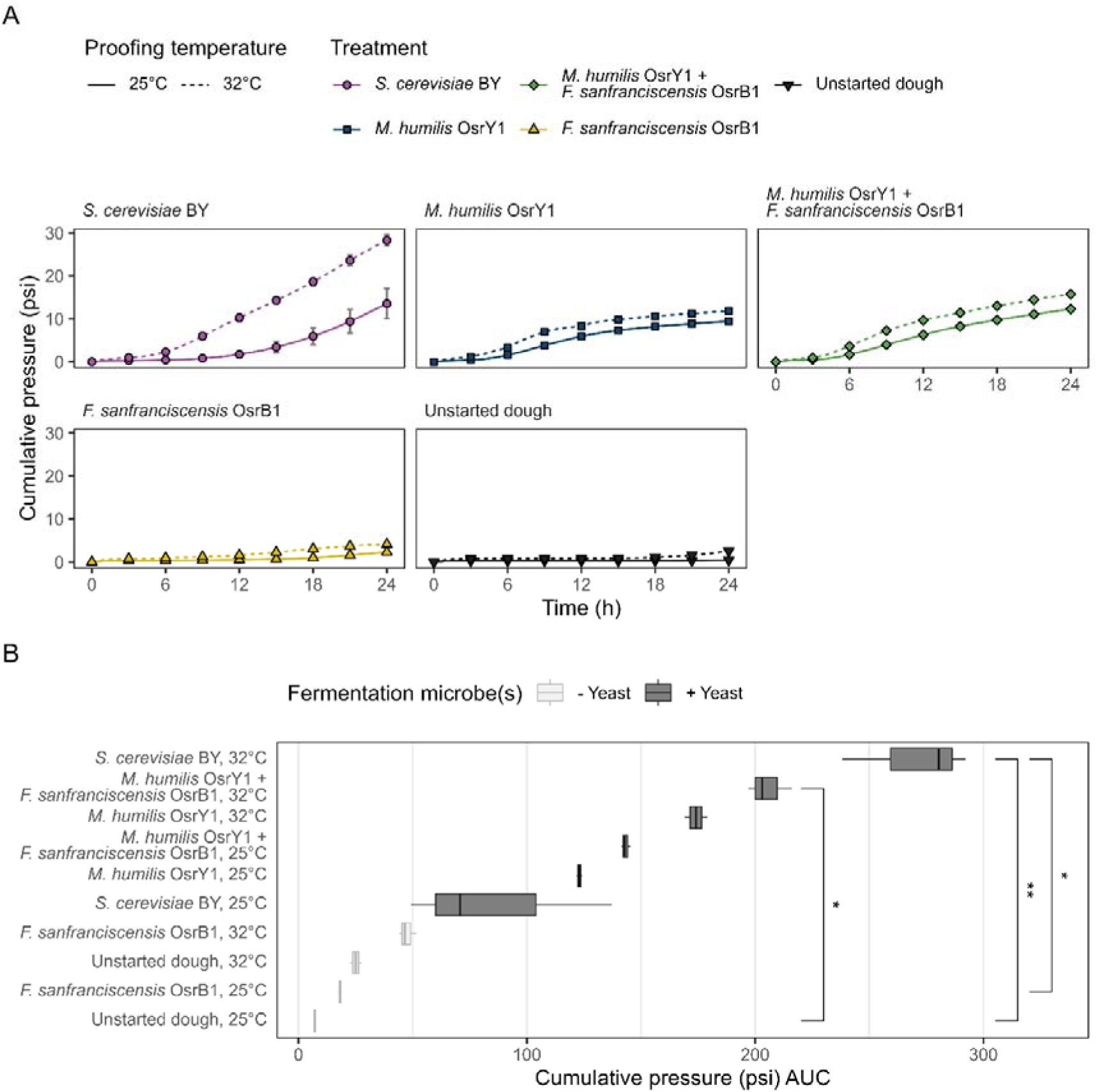
At 25 °C, the leavening ability of *M. humilis* OsrY1 in mono- and co-culture is comparable to that of baker’s yeast. **A** Lines indicate cumulative dough pressure output. **B** Boxplots show each treatment-temperature group AUC value. Mean (*n* = 3) values shown, error bars indicate SE. * *p* < 0.05, ** *p* < 0.01.

### FODMAP content

Changes in concentrations of key dough FODMAPs (fructans, mannitol, excess fructose) over time under different fermentation conditions were measured.

Fructan was the main FODMAP detected (89.6% of total measured FODMAP content), with mannitol and excess fructose being found in minor proportions (5.3% and 5.1%, respectively). Over 24 h dough fermentation, *M. humilis* OsrY1 was capable of fructan degradation (and lowering of total FODMAP content) to the same extent as baker’s yeast (**Figure 4**). By 24 h, all three yeasted treatments lowered fructan levels to well below the 0.3 g per serve threshold required to classify a food as low in oligosaccharides.

**Figure 4.**
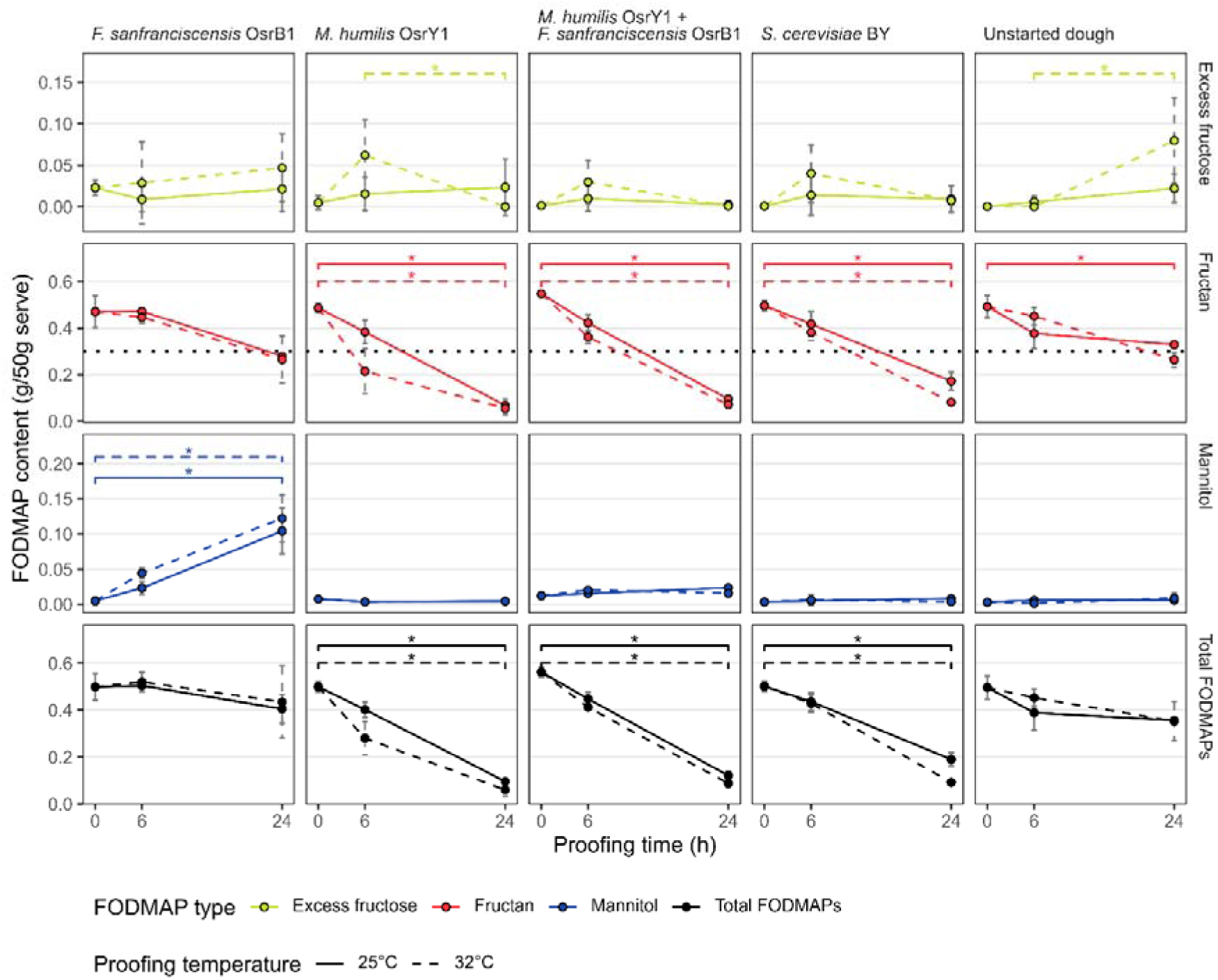
Reduction of fructan levels to below the low-FODMAP threshold is achieved when a fermenting yeast is present and the dough is fermented for 6 hours or longer. The dotted line indicates low-FODMAP threshold for oligosaccharides in cereal products as described by Varney et al. [3]. Points indicate mean (*n* = 3) total FODMAP content per 50 g serving ± SD. * *p* < 0.05.

The increase in mannitol concentration in the dough over time was only significant for dough fermented solely by *F. sanfranciscensis* OsrB1. By the 24 h timepoint, mannitol consisted of approx. 27% total FODMAP content in *F. sanfranciscensis* OsrB1 dough fermented at both 25 and 32°C. Furthermore, mannitol levels were negligible in all other treatments, including the dough co-cultured with *M. humilis* OsrY1.

In yeasted doughs fermented at 32°C, excess fructose tended to accumulate around the 6 h timepoint and deplete as fermentation continued. For *M. humilis* OsrY1, this depletion was statistically significant (*p* = 0.04291). In general, dough monosaccharides followed an ‘accumulate-deplete’ pattern in yeasted doughs (**Figure S2**). Available fructose and glucose levels were not linked to dough α-amylase content, which did not change in any inoculated doughs (**Figure S1**), so either microbial enzymes or other forms of endogenous amylases were likely to have been responsible.

At each time point, fermentation temperature demonstrated no significant effect on total FODMAP content. However, for specific treatment groups, effects related to individual FODMAPs were observed. At both 6 and 24 h fermentation, total FODMAP levels in *M. humilis* OsrY1 mono-culture doughs fermented at 32°C were lower than those at 25°C (*p* = 0.04953). *M. humilis* OsrY1 dough proofed for 6 h at 32°C was the only treatment with fructan levels below the 0.3 g per serve threshold at this time point. Fructan degradation by baker’s yeast was slightly improved at 32°C compared to 25°C at the 24 h time point (*p* = 0.04953), and doughs containing *F. sanfranciscensis* OsrB1 showed significantly higher mannitol levels at 6 h when fermented at 32°C. In general, temperature-based comparisons were only marginally significant at the α = 0.05 threshold. There were also no significant differences in excess fructose content according to proofing temperature.

### Crumb structure quality parameters

To ensure that any low-FODMAP bread produced was sufficiently leavened and had an optimal crumb structure, various loaf quality parameters were measured.

An interplay between fermentation microbe, temperature, and extent was observed in the degree of leavening of the different breads. In the early stages of fermentation (2 h), only loaf heights of yeasted treatments proofed at 32°C approached acceptable values. The baker’s yeast loaves were significantly taller at 2 and 6 h fermentation at 32 than at 25°C. For doughs baked at later fermentation stages, the distinction between yeasted and non-yeasted doughs was more evident at 25 than 32°C (**Figure 5A**). The yeasted doughs fermented at 32°C underwent over-proving (collapse of dough gluten structure resulting in reduced loaf height). This effect was most pronounced for the baker’s yeast bread, whose height decreased by approx. 10 mm by 24 h compared to 6 h. Bread is considered appropriately leavened when composed of approximately 25% ‘void’ or gas pockets (**Figure 5B**). The void fraction of the *M. humilis* OsrY1 mono- and co-culture loaves was not significantly different from the baker’s yeast loaves under any conditions, nor did the mono- and co-culture loaf void fractions differ from each other.

**Figure 5.**
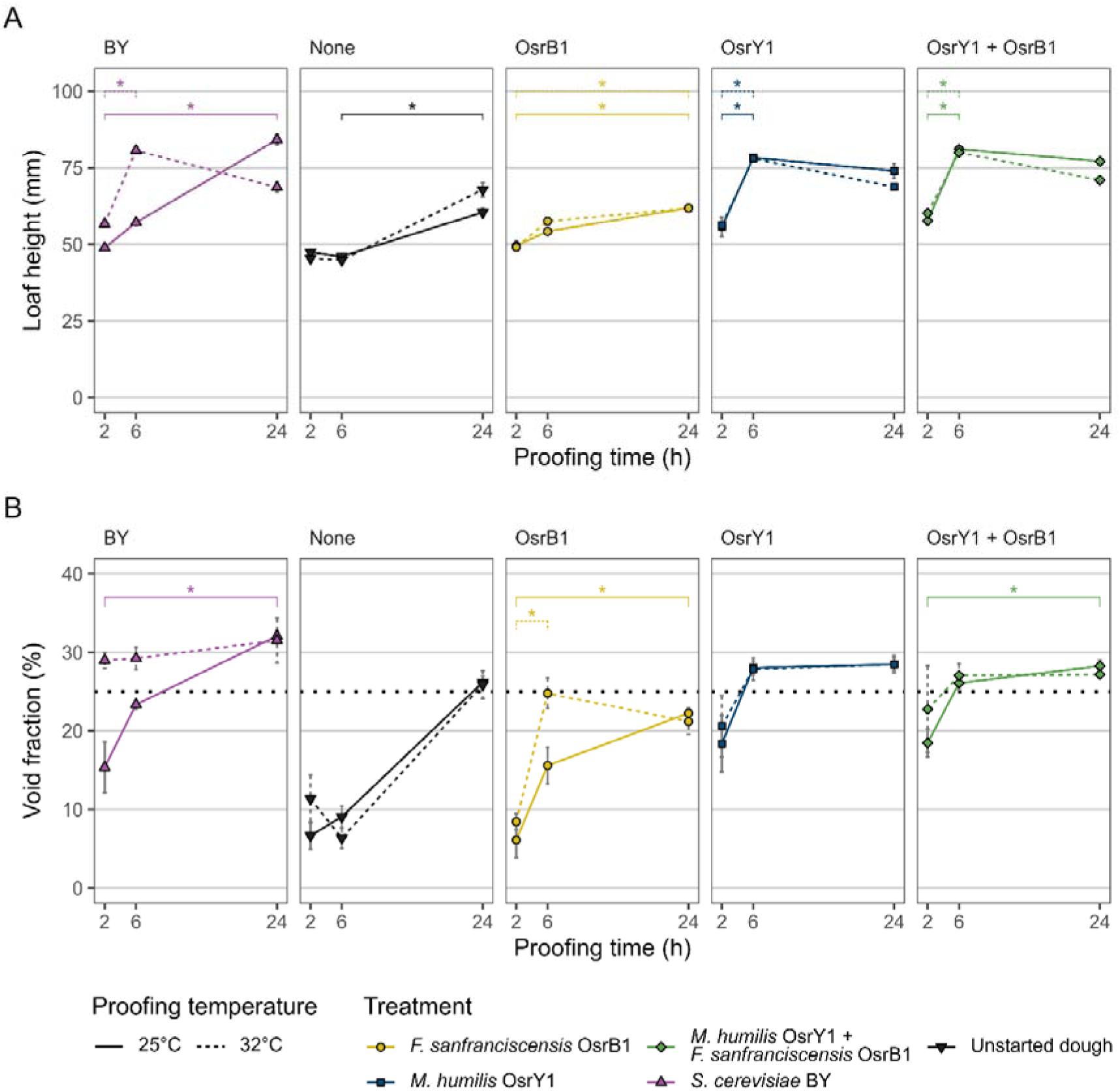
During dough fermentation, higher temperatures are associated with faster leavening, but cause more severe over-proofing. **A** Loaf height, **B** percentage of loaf cross section composed of ‘void’ or gas pockets (well-leavened bread exceeds 25% void fraction, indicated by the black dotted line). Points and error bars indicate mean (*n* = 3) ± SE. * *p* < 0.05.

The size and variation in size of the gas pockets is also an important crumb structure parameter. The average gas pocket size increases with fermentation duration (**Figure S4A**). A less uniform crumb structure was more common in bread made from dough proofed at 25°C, and large gas pockets only appeared in the 32°C breads once over-proved (**Figure S4B**).

### Bread sensory analysis

Finally, to confirm that low-FODMAP bread produced with sourdough co-cultures would be palatable, we conducted a sensory panel. Un-trained participants were asked to rate various visual, taste, and aroma aspects of each of the four samples on 7-point scales with defined extremes. **Figure 6** shows the distribution and mean scores given to each sample in response to the survey.

**Figure 6.**
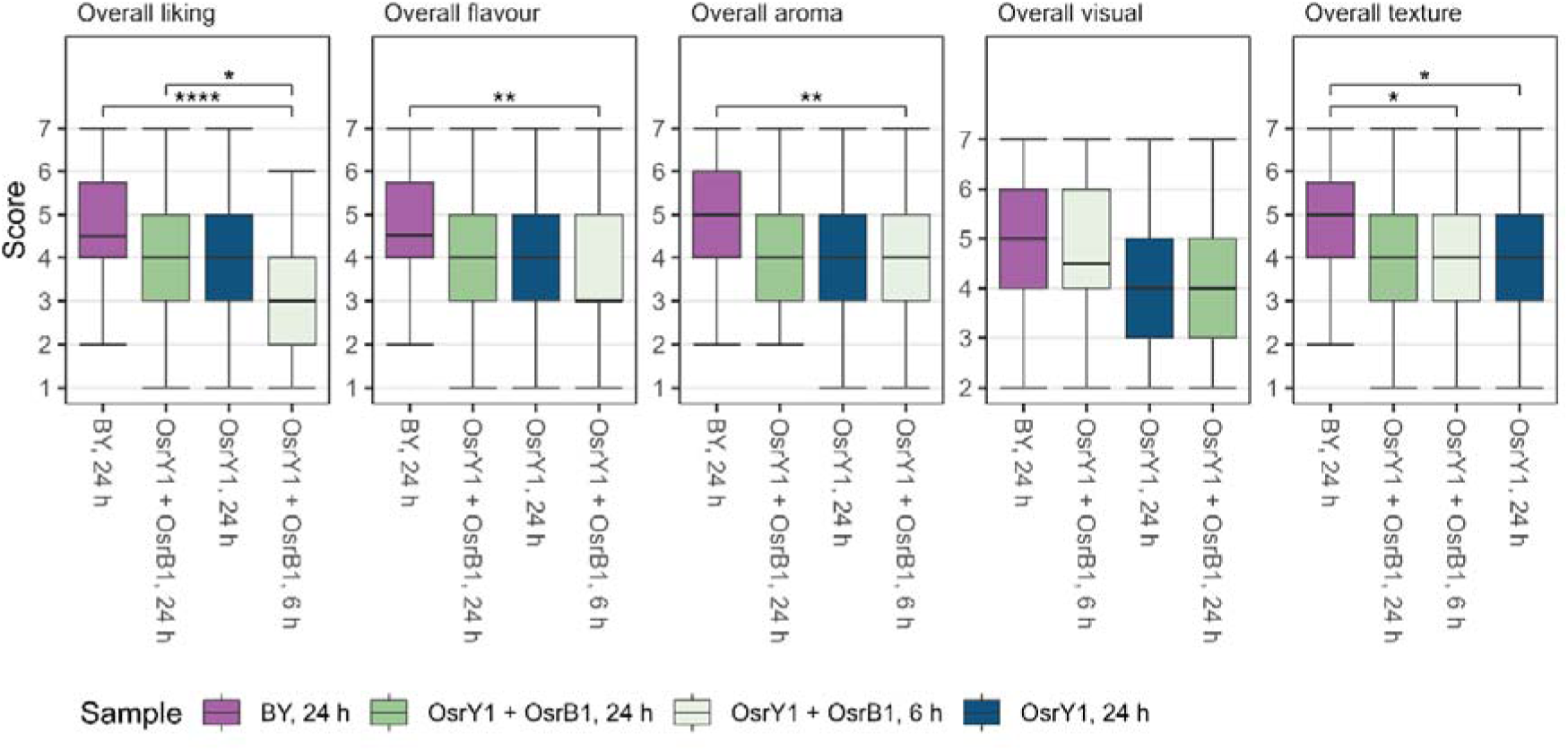
While standard bread is generally preferred, fermenting sourdough co-culture bread for longer improves overall liking and distinctive sourdough flavours and aromas. Un-trained study participants (*n* = 56) scored samples on each attribute on a 7-point scale with defined extremes (1 = dislike, 7 = like). *p* < 0.05, ** *p* < 0.01, **** *p* < 0.0001.

Sensory participants gave the highest overall liking scores to baker’s yeast bread fermented for 24 h at 25°C, and this was also reflected in their scores for overall flavour, aroma, and texture (**Figure 6**). The baker’s yeast sample only scored significantly higher for overall liking, flavour, and visual than the 6 h bread co-fermented with *M. humilis* and *F. sanfranciscensis*, not the two 24 h breads fermented with *M. humilis*.

Between the two breads fermented with sourdough co-culture, panellists tended to prefer the 24 h bread, which received higher scores for sour/vinegar and fruity aromas and flavours. The 24 h bread also had a lower total FODMAP content than the 6 h bread (**Figure 4**). The aroma complexity of the 24 h co-culture bread was also rated significantly higher than the 6 h bread. Both sour/vinegar attributes were scored higher for the 24 h co-culture bread than the *M. humilis* mono-culture bread although they did not differ in terms of dough pH (**Table S1**). Fruity aromas and flavours were also scored as significantly stronger in the co-culture 24 h bread than the 6 h bread.

Of the three 24 h breads, the two containing *M. humilis* were rated as being significantly darker in colour than the baker’s yeast bread, and overall visual scores were lower for these two samples than the baker’s yeast and 6 h co-culture breads (**Figure S3)**. Positive perceptions of bread texture may have been inversely related to dough pH (**Table S1).**

## Discussion

This study isolated and identified yeasts and bacteria in sourdough starters and optimised the use of a selected sourdough-derived co-culture in the production of low-FODMAP white wheat bread. Variation in bread dough fermentation process parameters facilitates leavening and FODMAP degradation to differing extents depending on the fermentation microbe(s) used.

Globally, 80 species of yeast and 82 species of lactobacilli have been identified in multiple instances in sourdough starters [17]. In accordance with the findings of our survey of sourdough starters, the two most common yeast species found in sourdough starters are *Saccharomyces cerevisiae* and *Maudiozyma humilis* [31, 32]. *M. humilis* and *F. sanfranciscensis* have previously been found to frequently co-occur in sourdough starters and contribute interdependently to the bread fermentation process [33]. This pairing occurred in three of the nine sourdough starters sampled in the present study.

Our quantification of fructans in white wheat dough aligns with previously published values, and confirms that they are the principal FODMAP found in bread dough [7, 10].

The inulinase activity of *Kluyveromyces marxianus* CBS6014 allows this yeast to effect rapid and near-complete degradation of fructans in whole wheat flour, whose fructan content is generally higher than that in white wheat flour [10, 24]. *M. humilis*, which does not produce inulinase or any other enzyme specific to dough oligosaccharides relies instead on invertase. Both our and other published studies of changes in fructan levels in dough fermented by invertase-producing yeasts like *M. humilis* and *S. cerevisiae* suggest that effectiveness of invertase-mediated fructan degradation is heavily dependent on fermentation duration [6, 13, 23]. Mannitol accumulated in the dough inoculated solely with *Fructilactobacillus sanfranciscensis*. This was expected, as this bacterium is known to use fructose in bread dough as an external electron acceptor to regenerate NADP, thereby producing mannitol [34]. However, it did not accumulate in the co-culture dough treatment. It is unlikely that *M. humilis* consumed the mannitol produced, as it is described as negative for growth or assimilation on mannitol [35]. The more likely explanation is that *M. humilis* consumed the available fructose before *F. sanfranciscensis* could convert it to mannitol, as in standard bread containing only baker’s yeast, fructose concentration decreases with extended fermentation duration [6]. This is further supported by depletion of fructose in yeasted doughs compared to the bacteria-only treatments (Figure S2).

Dough fermentation co-cultures must perform well from a technical standpoint as well as beneficially modulating bread FODMAP composition. Differences in fermentative performance according to dough fermentation temperature suggested that baker’s yeast is reliant on higher temperatures for optimal leavening and FODMAP reduction. This is likely to be a property of *S. cerevisiae* having a larger growth temperature range [36]. This difference was less pronounced for *M. humilis*, whose type strain does not grow at temperatures over 30°C [35]. Furthermore, we found that decreases in loaf height were worse at the higher dough fermentation temperature tested, although more specific dough gas retention capacity and porosity data were not able to be captured by the ANKOM gas pressure monitors used. Together, these results suggest that *M. humilis* is better suited to an extended, cooler dough fermentation regime targeted towards reducing FODMAP levels while maintaining bread quality. Only one pairing was tested in the present study, but Carbonetto et al. [37] found that co-culturing *M. humilis* with LAB strains affected loaf height in a strain combination-dependent manner. However, as an obligately heterofermentative bacterium, *Fructilactobacillus sanfranciscensis* is expected to affect loaf height in either a positive or neutral way [37], so it is possible that other sourdough-derived *F. sanfranciscensis* strains would interact with *M. humilis* in a similar way due to their physiology. Lactic acid produced by *F. sanfranciscensis* slightly weakens the dough gluten matrix by increasing its elasticity, necessitating dough manipulation technique adjustments to account for this [38]. Decreases in dough pH were also observed in yeast-only doughs (Table S1), although variability of these measures may have been affected by variables such as ambient humidity.

In addition to their loaf technical properties (i.e., leavening), fermentation microbes from sourdough contribute desirable aroma and flavour attributes to bread that consumers find appealing. Non-conventional yeast species are known to generate particular aromas in bread, especially fruity ones [39]. Our panellists only detected these in breads co-fermented with *F. sanfranciscensis*, suggesting that organic acids were the main volatile metabolites being detected. More generally, our study found that consumers preferred baker’s yeast bread, which echoes previous findings from similar trials [40]. However, while panellists’ preferences for the baker’s yeast and co-culture breads fermented for 24 h at 25°C were similar, significant differences between 6 and 24 h sourdough co-culture breads were detected. In terms of overall liking, colour, sour/vinegar aromas and flavours, fruity aromas and flavours, and aroma complexity, the 24 h sourdough co-culture breads scored higher. This demonstrates the potential sensory value in extending dough fermentations. Many sourdough-associated volatile aroma compounds evaporate before or during the baking process, however [41], which may explain why the sourdough microbe breads did not receive the highest scores for aroma complexity.

The research presented here builds on the small but growing body of literature focusing on Australian sourdough starters. Our results show that increasing fermentation temperature slightly improves the extent of fructan degradation by invertase-producing yeasts. However, this effect is only relevant to the early stages of dough fermentation (< 6 h). Likewise, higher fermentation temperatures lead to slightly improved leavening in yeasted doughs, but this leads to over-proving in extended dough fermentations. This effect was less pronounced for *M. humilis*, whose gas production pattern was slower, steadier, and less affected by proofing temperature increases. Therefore, we recommend that a dough fermentation aimed at FODMAP reduction should feature an extended dough fermentation (> 6 h), ambient proofing temperature, and fermenting microbes which do not over-leaven bread in these conditions.

## Acknowledgements

We wish to thank bakers of Victoria who donated samples of their sourdough starters to us. For their time and effort, we wish to thank all bread sensory study participants. Julie Hanot assisted in the collection of sourdough starter samples and yeast isolation, and her work is gratefully acknowledged.

## Human ethics and consent to participate

Every participant gave free and informed consent.

## Ethical approval

The sensory assessment performed in this study was approved by The University of Melbourne Human Research Ethics Committee (HREC 2024-30068-56367-3), in accordance with the Declaration of Helsinki.

## Author contribution declarations

Anna Elizabeth Wittwer and Jiaying Amanda Li conducted the experiments, analysed data and curated resources. Anna Elizabeth Wittwer wrote the first draft of the paper. Kate Howell provided oversight and supervision of the project. All authors reviewed and approved the manuscript.

