## Supplementary files for "Applying deconstructed sourdough communities and fermentation parameters to low-FODMAP wheat bread production"

Supplementary materials

Table S1 Changes in dough pH over time. Mean values ± SD shown.

| Proofing temperature | Proofing time | BY | OsrB1 | OsrY1 | OsrY1 + OsrB1 | None |
| --- | --- | --- | --- | --- | --- | --- |
| 25°C | 0 h | 5.86 ± 0.08 | 5.87 ± 0.05 | 5.87 ± 0.01 | 5.85 ± 0.04 | 5.9 ± 0.1 |
| 25°C | 2 h | 5.89 ± 0.07 | 5.9 ± 0.02 | 5.96 ± 0.03 | 5.88 ± 0.04 | 5.92 ± 0.08 |
| 25°C | 6 h | 5.94 ± 0.14 | 5.95 ± 0.08 | 5.91 ± 0.05 | 5.78 ± 0.03 | 5.94 ± 0.07 |
| 25°C | 24 h | 5.55 ± 0.04 | 5.32 ± 0.07 | 5.31 ± 0.1 | 5.24 ± 0.01 | 5.76 ± 0.12 |
| 32°C | 0 h | 5.86 ± 0.04 | 5.86 ± 0.02 | 5.87 ± 0.01 | 5.87 ± 0.03 | 5.86 ± 0.08 |
| 32°C | 2 h | 5.88 ± 0.06 | 5.87 ± 0.04 | 5.89 ± 0.02 | 5.86 ± 0.03 | 5.88 ± 0.05 |
| 32°C | 6 h | 5.8 ± 0.19 | 5.76 ± 0.11 | 5.68 ± 0.06 | 5.61 ± 0.04 | 5.88 ± 0.05 |
| 32°C | 24 h | 5.42 ± 0.04 | 5.03 ± 0.16 | 4.95 ± 0.43 | 5.11 ± 0.01 | 5.61 ± 0.12 |

Table S2 Changes in dough temperature over time. Mean values ± SD shown.

| Proofing temperature | Proofing time | BY | OsrB1 | OsrY1 | OsrY1 + OsrB1 | None |
| --- | --- | --- | --- | --- | --- | --- |
| 25°C | 0 h | 21.77 ± 0.77 | 22.01 ± 0.54 | 21.33 ± 0.28 | 22.03 ± 0.6 | 22.1 ± 0.81 |
| 25°C | 2 h | 23.84 ± 0.13 | 23.81 ± 0.14 | 23.51 ± 0.55 | 24.42 ± 0.05 | 23.64 ± 0.19 |
| 25°C | 6 h | 24 ± 0.22 | 24.16 ± 0.42 | 23.96 ± 0.1 | 24.37 ± 0.23 | 23.8 ± 0.18 |
| 25°C | 24 h | 24.17 ± 0.47 | 23.96 ± 0.39 | 24.1 ± 0.38 | 24.38 ± 0.21 | 23.86 ± 0.62 |
| 32°C | 0 h | 21.66 ± 0.86 | 21.54 ± 0.23 | 21.44 ± 0.16 | 21.92 ± 0.63 | 21.96 ± 1.06 |
| 32°C | 2 h | 28.07 ± 0.58 | 29.17 ± 0.61 | 28.74 ± 0.28 | 29.14 ± 1 | 28.23 ± 0.4 |
| 32°C | 6 h | 29.07 ± 0.99 | 29.11 ± 0.53 | 29.18 ± 0.53 | 29.94 ± 0.08 | 28.18 ± 0.95 |
| 32°C | 24 h | 28.38 ± 0.96 | 28.66 ± 0.07 | 28.08 ± 0.43 | 29.2 ± 0.21 | 27.96 ± 0.49 |

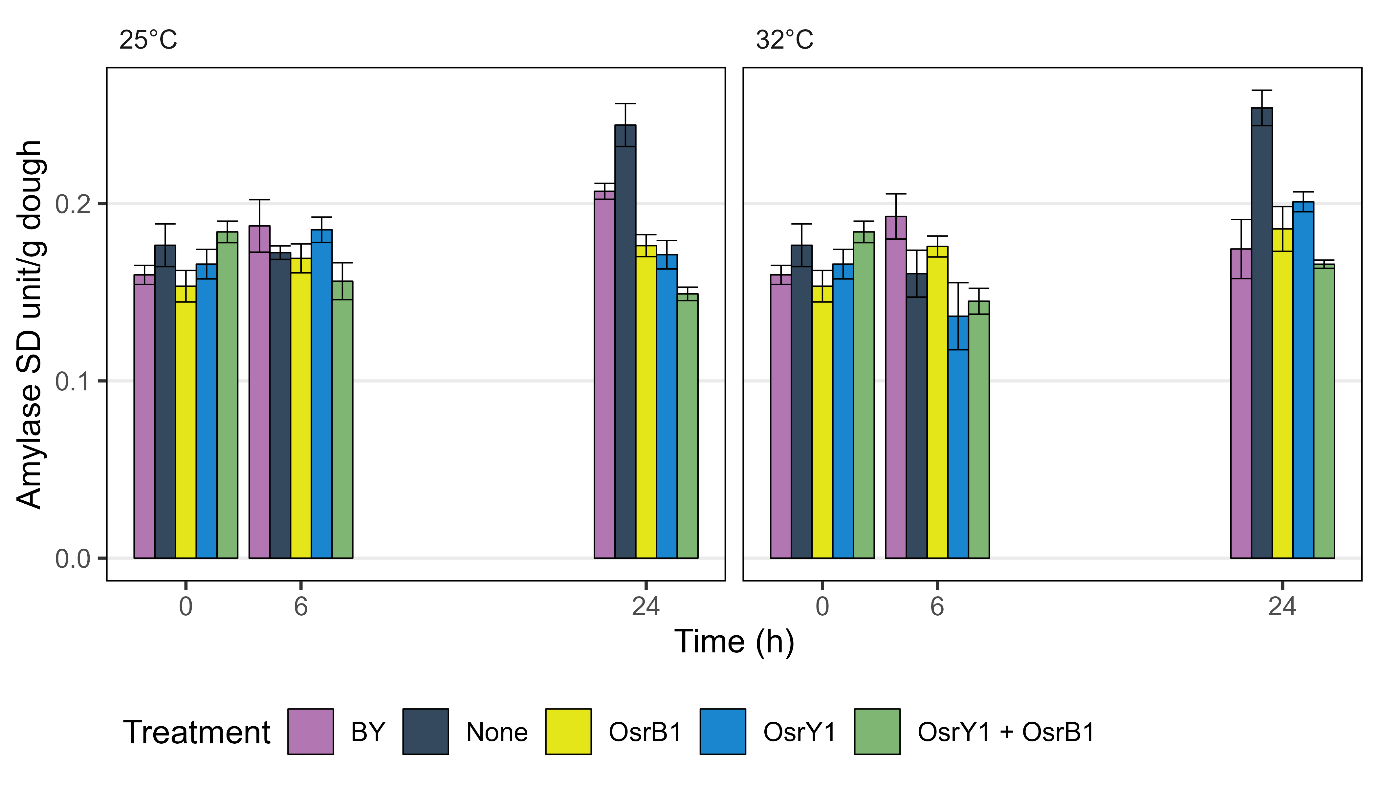

Figure S1 Dough α-amylase levels over time. Bars indicate mean values (*n* = 3) ± SE.

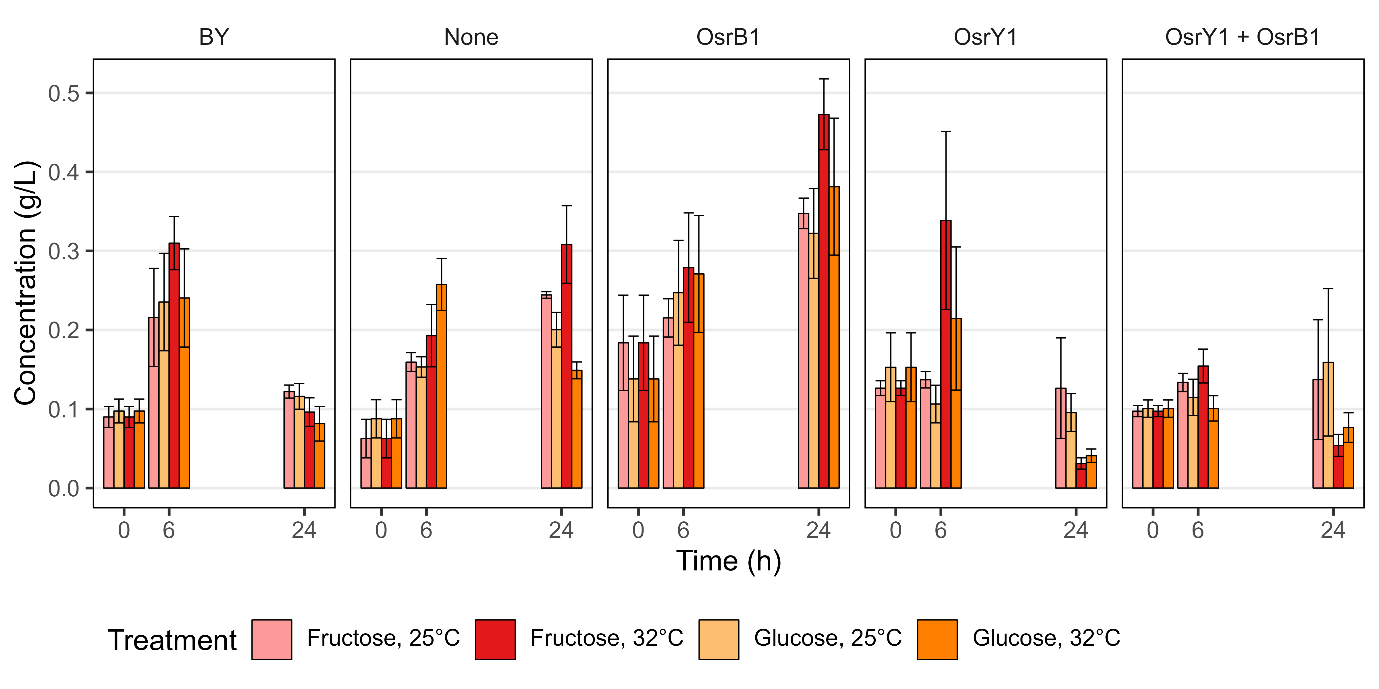

Figure S2 Concentrations of glucose and fructose in dough fermented at different temperatures. Bars indicate mean values (*n* = 3) ± SE.

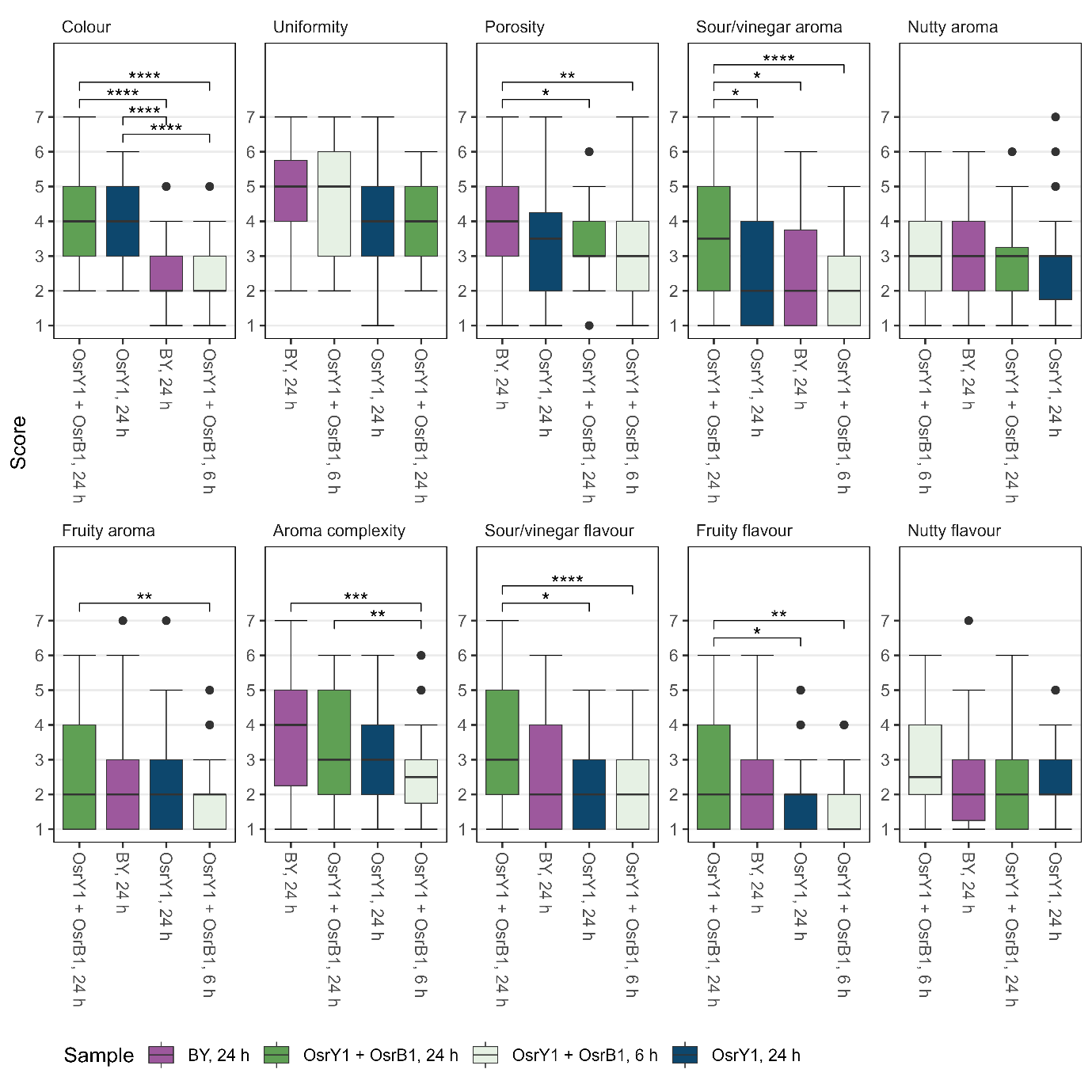

Figure S3 Supplementary sensory panel data. Un-trained study participants (*n* = 56) scored samples on each attribute on a 7-point scale with defined extremes (1 = dislike, 7 = like). *p* < 0.05, ** *p* < 0.01, *** *p* < 0.001, **** *p* < 0.0001.

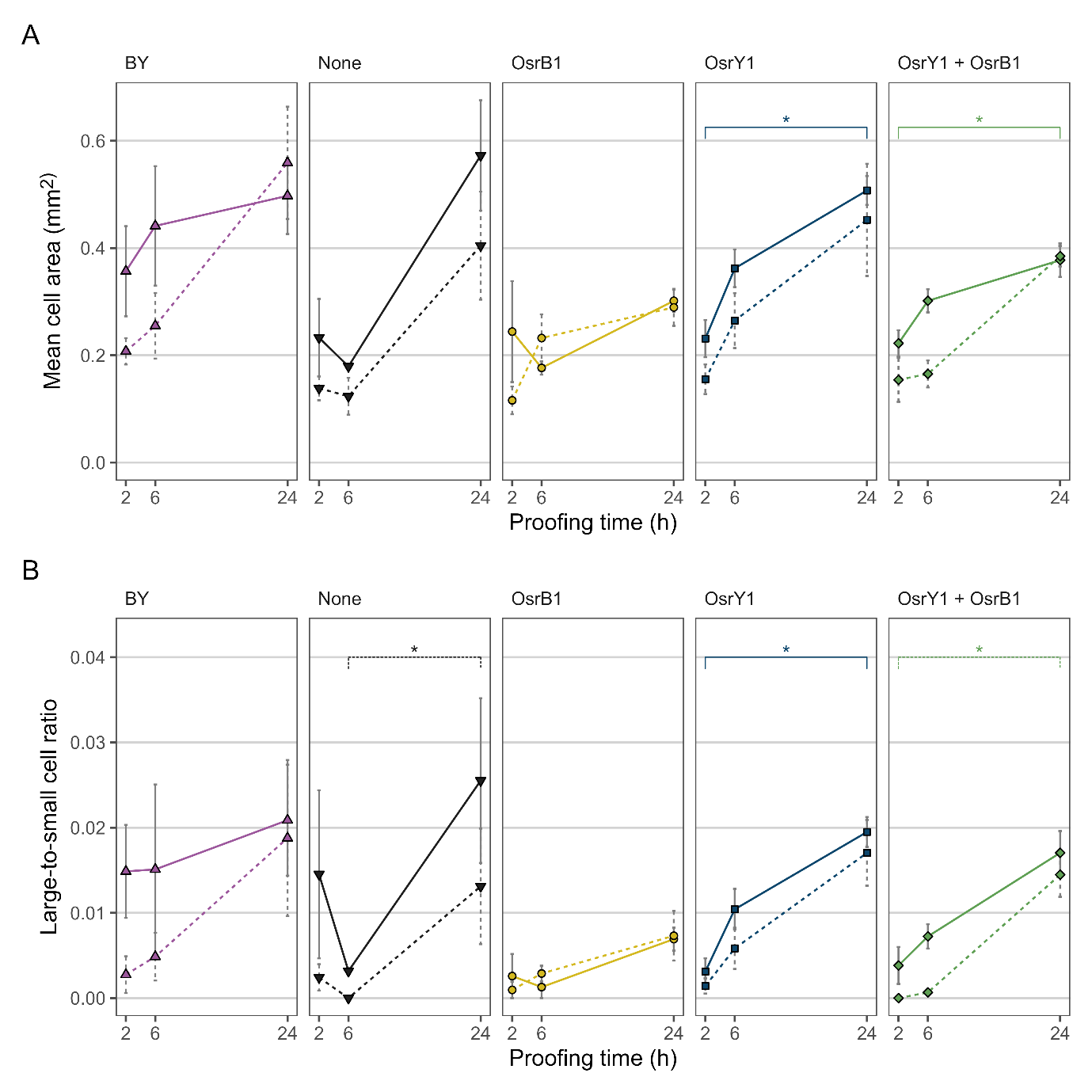

Figure S4 Supplementary loaf crumb structure parameters. **A** Average gas pocket size, and **B** the ratio of large to small gas pockets. Points and error bars indicate mean (*n* = 3) ± SE. * *p* < 0.05.
